# The cargo receptor SURF4 promotes the efficient cellular secretion of PCSK9

**DOI:** 10.1101/336016

**Authors:** Brian T. Emmer, Geoffrey G. Hesketh, Emilee Kotnik, Vi T. Tang, Paul J. Lascuna, Jie Xiang, Anne-Claude Gingras, Xiao-Wei Chen, David Ginsburg

## Abstract

Proprotein convertase subtilisin/kexin type 9 (PCSK9) is a secreted protein that plays an important role in regulating plasma cholesterol and cardiovascular disease risk. PCSK9 secretion uniquely depends on the cytoplasmic COPII protein SEC24A, suggesting the presence of a transmembrane ER cargo receptor mediating this interaction. Here, we report a novel approach that combines proximity-dependent biotinylation and proteomics together with genome-scale CRISPR screening to identify proteins that facilitate the efficient secretion of PCSK9 heterologously expressed in HEK293T cells. We first identified 35 candidate proteins that were labeled by BirA* fusions to PCSK9 and either COPII component SAR1A or SAR1B. We then performed genome-scale pooled CRISPR mutagenesis to identify genes whose perturbation resulted in intracellular accumulation of PCSK9-eGFP but not the control A1AT-mCherry. The 4 most enriched sgRNAs in this screen all targeted *SURF4*, a homologue of the yeast endoplasmic reticulum (ER) cargo receptor Erv29p and the only candidate also identified by proximity-dependent biotinylation. The functional contribution of *SURF4* to PCSK9 secretion was confirmed with multiple independent *SURF4*-targeting sgRNAs, clonal *SURF4*-deficient cell lines, and functional rescue with *SURF4* cDNA. Compatible with a function of *SURF4* as a cargo receptor for PCSK9, fluorescence microscopy localized *SURF4* to the early secretory pathway, coimmunoprecipitation revealed a physical interaction between *SURF4* and PCSK9, and *SURF4* deletion resulted in decreased extracellular secretion of PCSK9 and PCSK9 accumulation in the ER. Taken together, these findings support a model in which *SURF4* functions as an ER cargo receptor for the efficient cellular secretion of PCSK9.

## INTRODUCTION

PCSK9 is a proprotein convertase that acts as a negative regulator of the LDL receptor^1^. PCSK9 is synthesized primarily in hepatocytes and secreted into the bloodstream. Circulating PCSK9 binds to the LDL receptor and diverts it to lysosomes for degradation, thereby leading to decreased LDL receptor abundance at the hepatocyte cell surface, decreased LDL clearance, and hypercholesterolemia. PCSK9 was originally implicated in cardiovascular disease when human genetic studies identified gain-of-function PCSK9 mutations as a cause of familial hypercholesterolemia^2^. Subsequently, loss-of-function PCSK9 variants were associated with decreased plasma cholesterol and lowered lifetime incidence of cardiovascular disease^3,4^.Therapeutic inhibitors of PCSK9 have been recently developed that exhibit potent lipid-lowering effects and are associated with a reduction in cardiovascular events^5,6^.

A critical early sorting step for secreted proteins is their incorporation into membrane-bound vesicles that transport cargoes from the ER to the Golgi apparatus^7^. The formation of these vesicles is driven by coat protein complex II (COPII), which includes the SAR1 GTPase, heterodimers of SEC23/SEC24, and heterotetramers of SEC13/SEC31. Secreted cargoes are incorporated into COPII vesicles by two mechanisms. “Cargo capture” refers to concentrative, receptor-mediated, active sorting of selected cargoes, in contrast to “bulk flow”, by which cargoes enter COPII vesicles through passive diffusion. These mechanisms are not mutually exclusive, as cargoes may exhibit basal rates of secretion that are enhanced by receptor-mediated recruitment. It remains unclear to what extent protein recruitment into the secretory pathway is driven by selective cargo capture versus passive bulk flow^8^.

In yeast, the active sorting of secreted cargoes into COPII-coated vesicles is driven primarily by Sec24, which is encoded by a single, essential gene. In mammals, multiple paralogs of SEC24 exist, potentially accommodating a more diverse and regulated repertoire of cargoes. Genetic deficiency in the mouse for one of these paralogs, SEC24A, results in hypocholesterolemia due to reduced secretion of PCSK9 from hepatocytes^9^. This finding suggested an active receptor-mediated mechanism for PCSK9 recruitment into COPII vesicles. A direct physical interaction between SEC24A and PCSK9, however, is implausible since SEC24A localizes to the cytoplasmic side of the ER membrane and PCSK9 to the luminal side, with neither possessing a transmembrane domain. This topology instead implies the presence of an ER cargo receptor, a transmembrane protein that could serve as an intermediary between the COPII coat and luminal PCSK9.

Although COPII-dependent ER cargo receptors were first identified in yeast nearly two decades ago, few examples of similar cargo receptor interactions have been reported for mammalian secreted proteins^8^. Previous investigation of the ER cargo receptor LMAN1 demonstrated no specificity for SEC24A over other SEC24 paralogs, making this unlikely to serve as a PCSK9 cargo receptor^10^. Earlier analyses of PCSK9-interacting proteins^11-13^did not identify a clear receptor mediating PCSK9 secretion. Here, we developed a novel strategy for ER cargo receptor identification that combines proximity-dependent biotinylation with CRISPR-mediated functional genomic screening. This approach led to the identification of the ER cargo receptor *SURF4* as a primary mediator of PCSK9 secretion in HEK293T cells.

## RESULTS

### Identification of candidate PCSK9 cargo receptors by proximity-dependent biotinylation

To identify PCSK9-interacting proteins, we first engineered cells expressing a fusion of PCSK9 and a mutant biotin ligase, *E. coli* BirA*(R118G), that catalyzes proximity-dependent biotinylation of primary amines on neighboring proteins within an estimated ∼10 nm radius^14,15^, effectively converting transient interactions into covalent modification (Figure 1A). The high affinity of the biotin-streptavidin interaction in turn allows for stringent detergent and high salt conditions during purification. Quantitative mass spectrometry of streptavidin-purified interacting proteins from cells expressing PCSK9-BirA* identified 162 prey proteins that were specifically labeled (Bayesian FDR ≤1%) by PCSK9-BirA* relative to control bait proteins (Supplemental Table 1).

**Figure 1:**
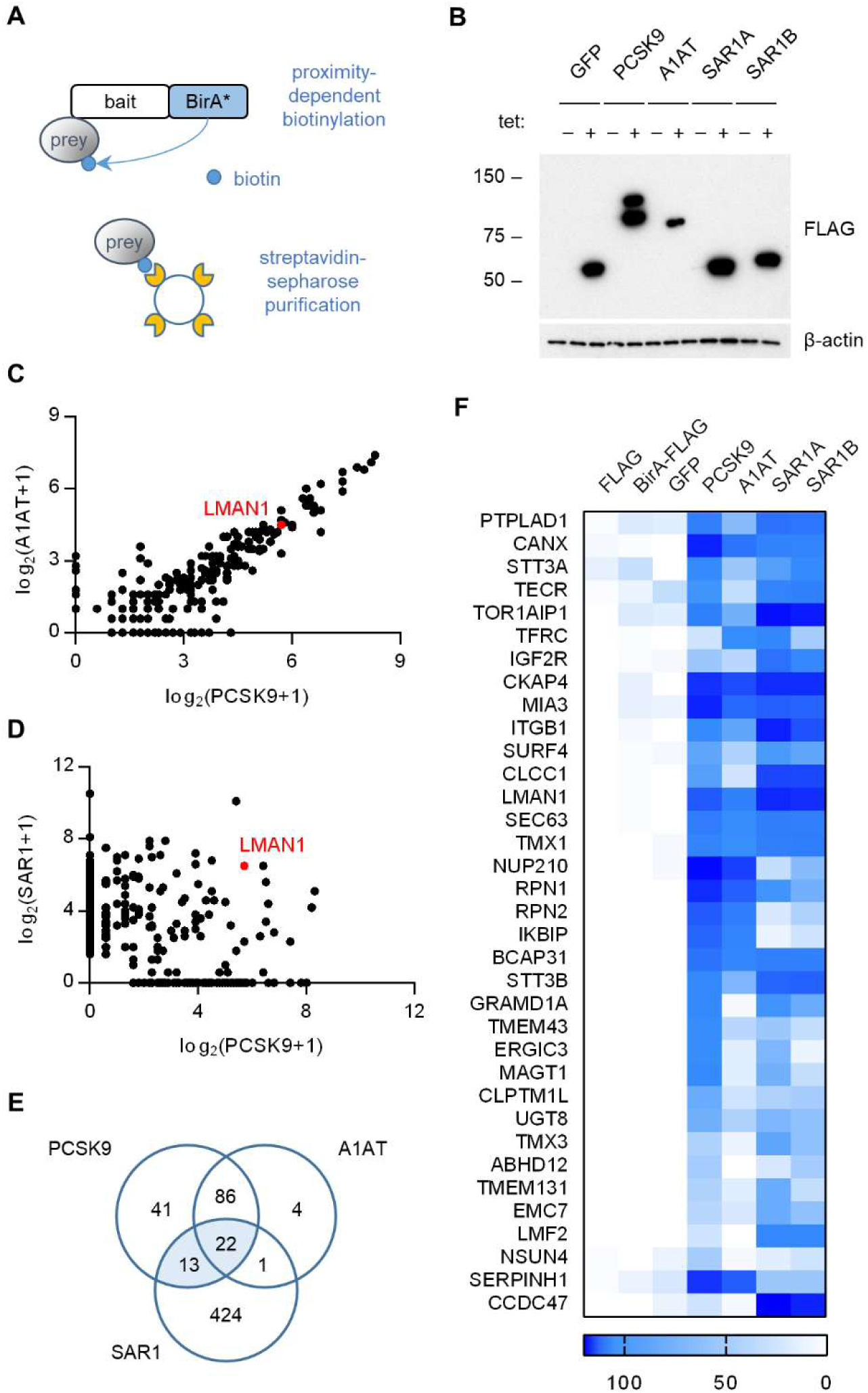
Proximity-dependent biotinylation with a PCSK9-BirA* fusion. **(A)** Proximity detection by mass spectrometry of streptavidin-purified prey proteins biotinylated by a fusion of BirA* to a bait protein of interest. (B) Immunoblotting of lysates of cells expressing various BirA*-fusion proteins. (C) Spectral counts of prey proteins identified from lysates of cells expressing PCSK9-BirA* relative to A1AT-BirA* (BFDR≤0.01 for either bait). (D) Spectral counts of prey proteins purified from lysates of cells expressing PCSK9-BirA* relative to the maximum spectral count from lysates of cells expressing either SAR1A-BirA* or SAR1B-BirA*. (E) Venn diagram of identified prey proteins from lysates of cells expressing BirA* fusions with PCSK9, A1AT, or the maximum for either Sar1A or Sar1B. (F) Heat map of spectral counts for candidate proteins demonstrating interaction with both PCSK9-BirA* and either SAR1A-BirA* or SAR1B-BirA*.

To refine the candidate list of PCSK9-interacting proteins, we next analyzed cells expressing a fusion of BirA* with a control secreted protein, alpha-1 antitrypsin (A1AT). The interactome of A1AT showed substantial overlap with that of PCSK9 (108/162 proteins, Figure 1C). The A1AT cargo receptor LMAN1 was similarly labeled by both PCSK9-BirA* and A1AT-BirA*, suggesting that the restricted environment of the COPII vesicle may lead to nonspecific labeling of adjacent cargo receptors. We next compared the interactome of PCSK9 to that of SAR1A and SAR1B (Figure 1D), COPII proteins that localize to the cytoplasmic surface of budding COPII vesicles, identifying a total of 35 candidate ER cargo receptors interacting with both PCSK9 and either SAR1A or SAR1B (Figure 1E-F, Supplemental Table 1). The majority of these candidates were annotated as integral membrane proteins (32/35, p = 3×10^-16^) with localization in the ER (24/35, p = 1.6×10^-18^), as would be expected for an ER cargo receptor (Supplemental Table 1).

### A genome-scale CRISPR screen identifies *SURF4* as a putative ER cargo receptor for PCSK9

We next developed a functional screen to identify genes involved in PCSK9 secretion (Figure 2A). We reasoned that mutants with reduced PCSK9 exit from the ER would accumulate intracellular PCSK9, and that fusion of PCSK9 to eGFP would allow for a cell-autonomous, scalable, and selectable readout of PCSK9 accumulation. We generated clonal HEK293T cell lines stably co-expressing both a PCSK9-eGFP fusion and, as an internal control, alpha-1 antitrypsin fused with mCherry. Immunoblotting verified the efficient secretion of both fusion proteins from clonal reporter cell lines (Supplemental Figure 1).

Treatment of these reporter cells with brefeldin A, a pharmacologic inhibitor of COPII function, resulted in intracellular accumulation of both PCSK9-eGFP and A1AT-mCherry (Figure 2B), consistent with ER exit and secretion of both fusion proteins through the COPII pathway.CRISPR-mediated inhibition of the ER cargo receptor for A1AT, LMAN1^16,17^, resulted in intracellular accumulation of A1AT-mCherry with no effect on PCSK9-eGFP (Figure 2B). To screen for specific modifiers of PCSK9 secretion, we next sought to identify single guide RNAs (sgRNAs) that would induce accumulation of PCSK9-eGFP with no change in A1AT-mCherry fluorescence. We mutagenized the PCSK9-eGFP-2A-A1AT-mCherry reporter cell line with the GeCKOv2 pooled library of 123,411 sgRNAs that includes 6 independent sgRNAs targeting nearly every coding gene in the human genome^18^(Figure 2A). Mutants with aberrant PCSK9-eGFP fluorescence but normal A1AT-mCherry fluorescence were then isolated by flow cytometry, with integrated lentiviral sgRNA sequences quantified by deep sequencing and analyzed for enrichment in PCSK9-eGFP high cells. The coverage and distribution of sgRNA sequencing reads demonstrated maintenance of library complexity and high reproducibility between biological replicates (Supplemental Figure 2).

**Figure 2:**
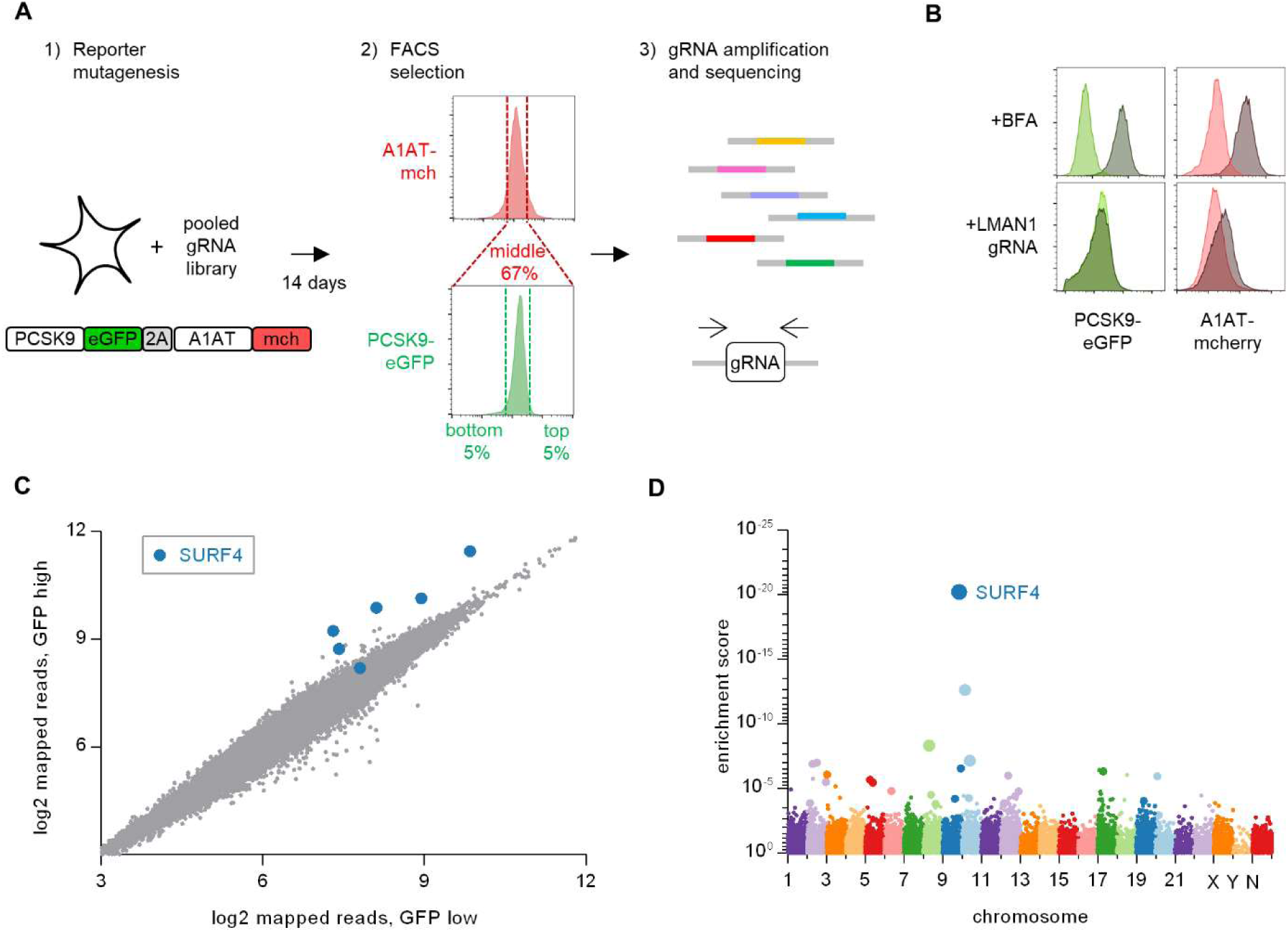
Whole genome CRISPR mutagenesis screen for PCSK9 secretion modifiers. (A) Strategy for whole genome screen. (B) Flow cytometry of reporter cells stably expressing PCSK9-eGFP-2A-A1AT-mCherry, treated with brefeldin A or a sgRNA targeting LMAN1. (C) Normalized abundance of each sgRNA in the library in eGFP high and eGFP low populations. (D) Gene-level enrichment scores. Diameter of the bubble for each gene is proportional to the number of unique sgRNAs significantly enriched in GFP high cells.

Strikingly, the 4 most enriched sgRNAs in the PCSK9-eGFP high cell population all targeted the same gene, *SURF4* (Figure 2C). The enrichment of *SURF4*-targeting sgRNA in eGFP-high cells was consistent across each of 4 biologic replicates and, after adjustment for multiple hypothesis testing, statistically significant for 5 of the 6 *SURF4*-targeting sgRNAs in the library (p <10^-13^– 10^-36^, Figure 3A). Gene-level analysis confirmed the strongest enrichment for *SURF4*-targeting sgRNA (Figure 2D). *SURF4* is a homologue of yeast Erv29, an ER cargo receptor that mediates the secretion of glycosylated pro-alpha-factor^19^. Comparison of candidate PCSK9 cargo receptors identified by either CRISPR functional screening (Supplemental Table 3) or proximity-dependent biotinylation (Figure 1F) demonstrated that *SURF4* was the only candidate identified by both approaches.

**Figure 3.**
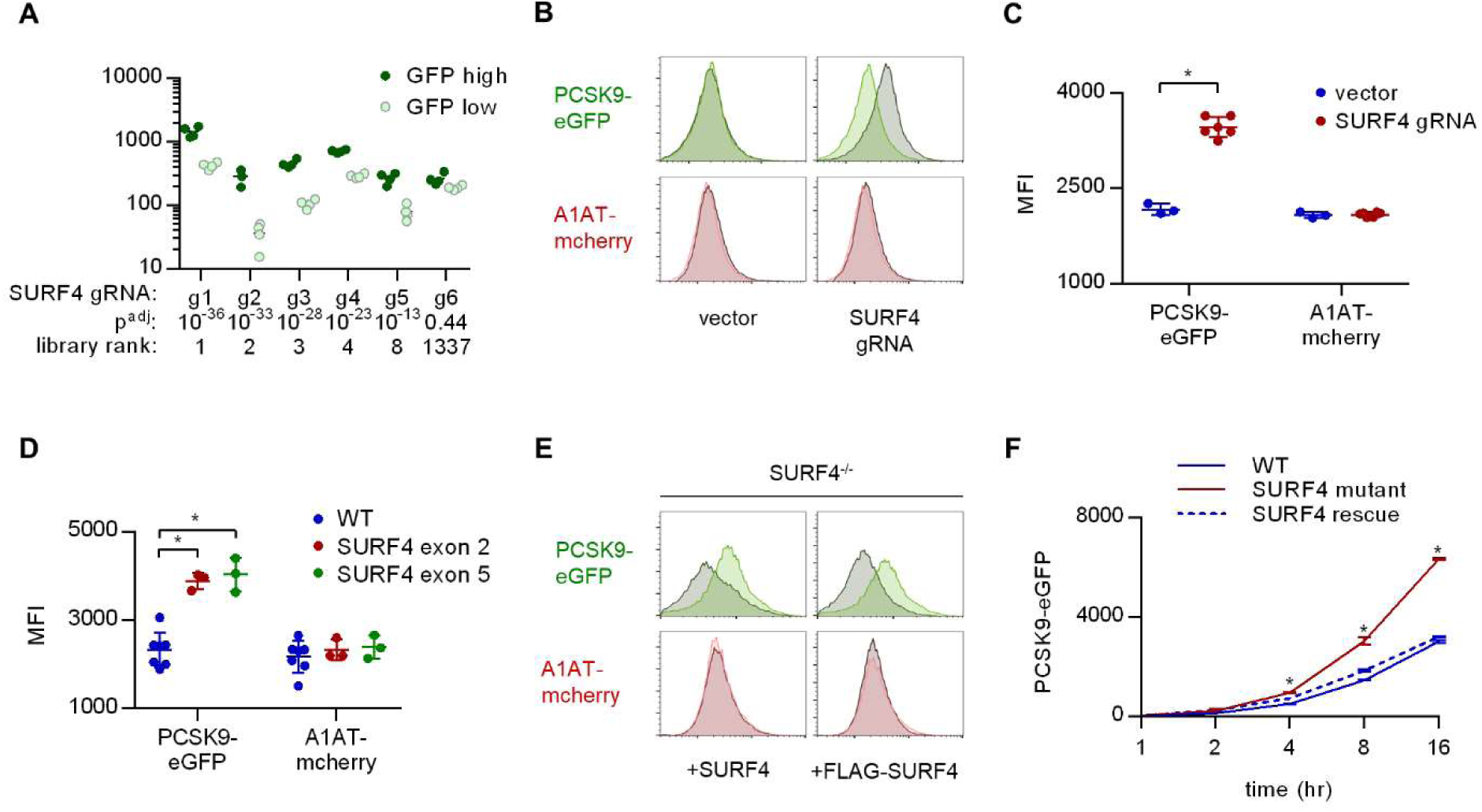
*SURF4* mutagenesis causes an accumulation of intracellular PCSK9-eGFP. (A) Individual sgRNA sequencing counts for *SURF4*-targeting sgRNA in eGFP high and eGFP low populations for each of 4 biologic replicates. (B) Validation by flow cytometry of PCSK9-eGFP and A1AT-mCherry fluorescence in reporter cells transfected with plasmids delivering Cas9 and *SURF4*-targeting sgRNA or empty vector. (C) Quantification of intracellular fluorescence for cells treated with vector or each of 6 unique *SURF4*-targeting sgRNAs. (D) Quantification of intracellular fluorescence for clonal cell lines each containing frameshift-causing indels at 2 different *SURF4* target sites. (E) Flow cytometry histograms for cells expressing PCSK9-eGFP-2A-A1AT-mCherry and deleted for *SURF4* with or without stable expression of a wild-type or FLAG-tagged *SURF4* cDNA. (F) Time course of intracellular accumulation of tetracycline-inducible PCSK9-eGFP on WT, *SURF4*-deficient, or *SURF4* rescue background.

### SURF4 deletion causes intracellular accumulation of PCSK9-eGFP in HEK293T cells

To validate the functional interaction of *SURF4* with PCSK9-eGFP, we generated plasmids encoding Cas9 and 6 independent *SURF4*-targeting sgRNAs (3 from the original screen and 3 additional unique sgRNAs). Reporter cells were transiently transfected with each of these 6 *SURF4*-targeting constructs and analyzed by FACS, with all 6 sgRNAs resulting in accumulation of intracellular PCSK9-eGFP fluorescence with no effect on intracellular A1AT-mCherry fluorescence (Figure 3B-C). Similarly, 6 clonal cell lines carrying sequence-verified indels in either *SURF4* exon 2 or exon 5 (Supplemental Figure 3) all exhibited specific PCSK9-eGFP accumulation with no effect on A1AT-mCherry (Figure 3D). This phenotype was rescued by stable expression of wild-type *SURF4* cDNA (Figure 3E). The intracellular accumulation of PCSK9 upon *SURF4* disruption was confirmed in an independently-derived HEK293T cell line carrying an inducible PCSK9-eGFP allele. Accumulation of PCSK9-eGFP in *SURF4* mutant cells relative to wild-type cells was detectable within 4 hr of induction (Figure 3F).

To exclude interaction of *SURF4* with the GFP portion of the fusion rather than PCSK9 itself, we also examined *SURF4*-dependence for the secretion of untagged PCSK9. Stable cell lines were generated by Flp/FRT-mediated knock-in of PCSK9 coding sequence into a tetracycline-inducible locus of HEK293T cells that were either wild-type or *SURF4*-deficient, the latter with or without stable integration of a wild-type *SURF4* cDNA expression construct. Consistent with the reduced secretion observed above for the PCSK9-eGFP fusion, untagged PCSK9 also exhibited intracellular accumulation in the absence of *SURF4* that was rescued by expression of a *SURF4* transgene (Figure 5B-C). Deletion of *SURF4* in these cells did not induce activation of ER stress markers (Supplemental Figure 4).

**Figure 4.**
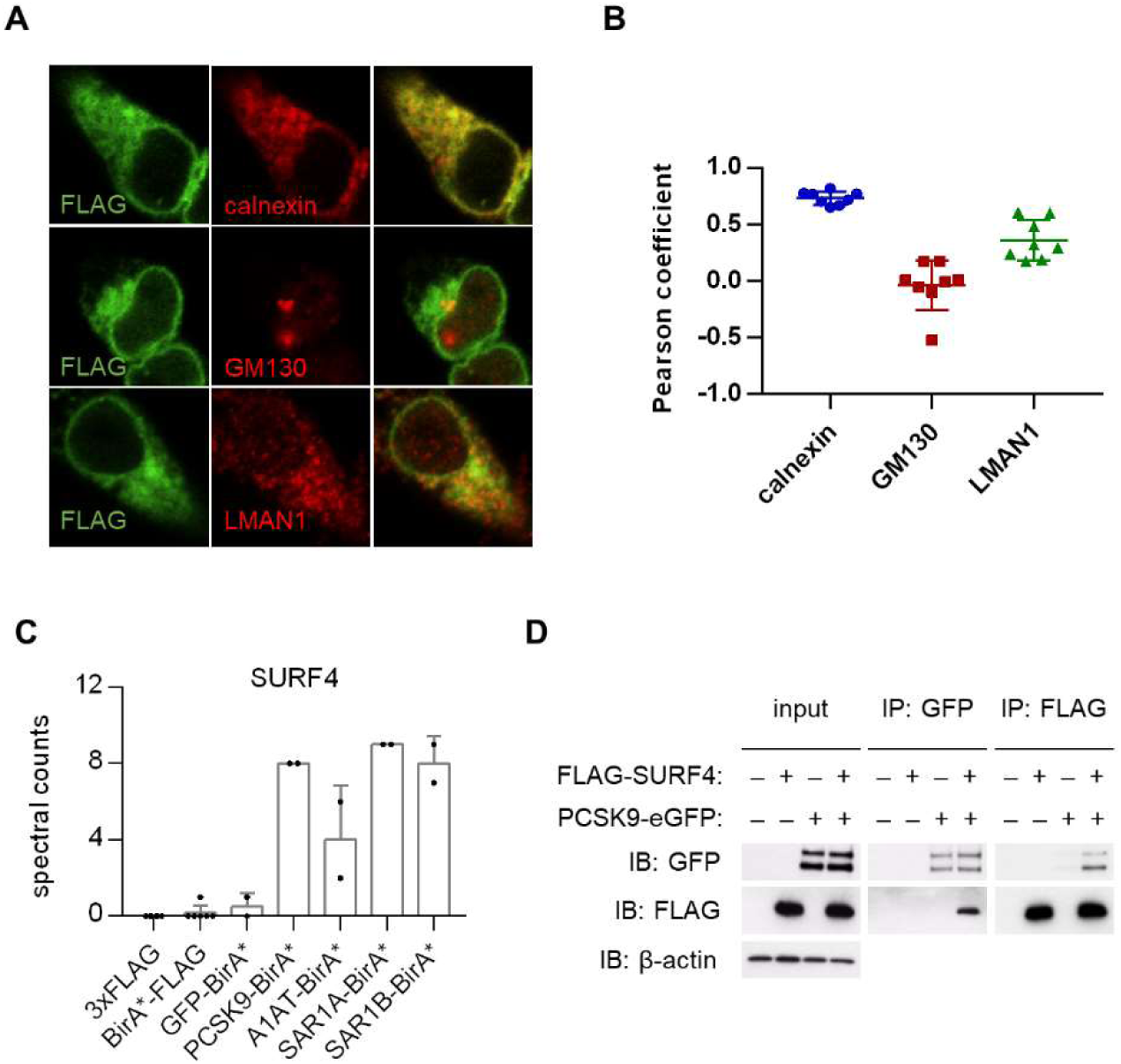
*SURF4* localizes to the early secretory pathway where it physically interacts with PCSK9. (A) Immunofluorescence of FLAG-*SURF4* together with markers of the ER (calnexin), ERGIC (LMAN1), and Golgi (GM130). (B) Quantification of colocalization. (C) Spectral counts for *SURF4* in streptavidin-purified eluates from cells expressing various BirA* fusion proteins. (D) Immunoprecipitations were performed using antibodies directed against FLAG or GFP from lysates of cells expressing FLAG-*SURF4*, PCSK9-eGFP, both, or neither.

**Figure 5.**
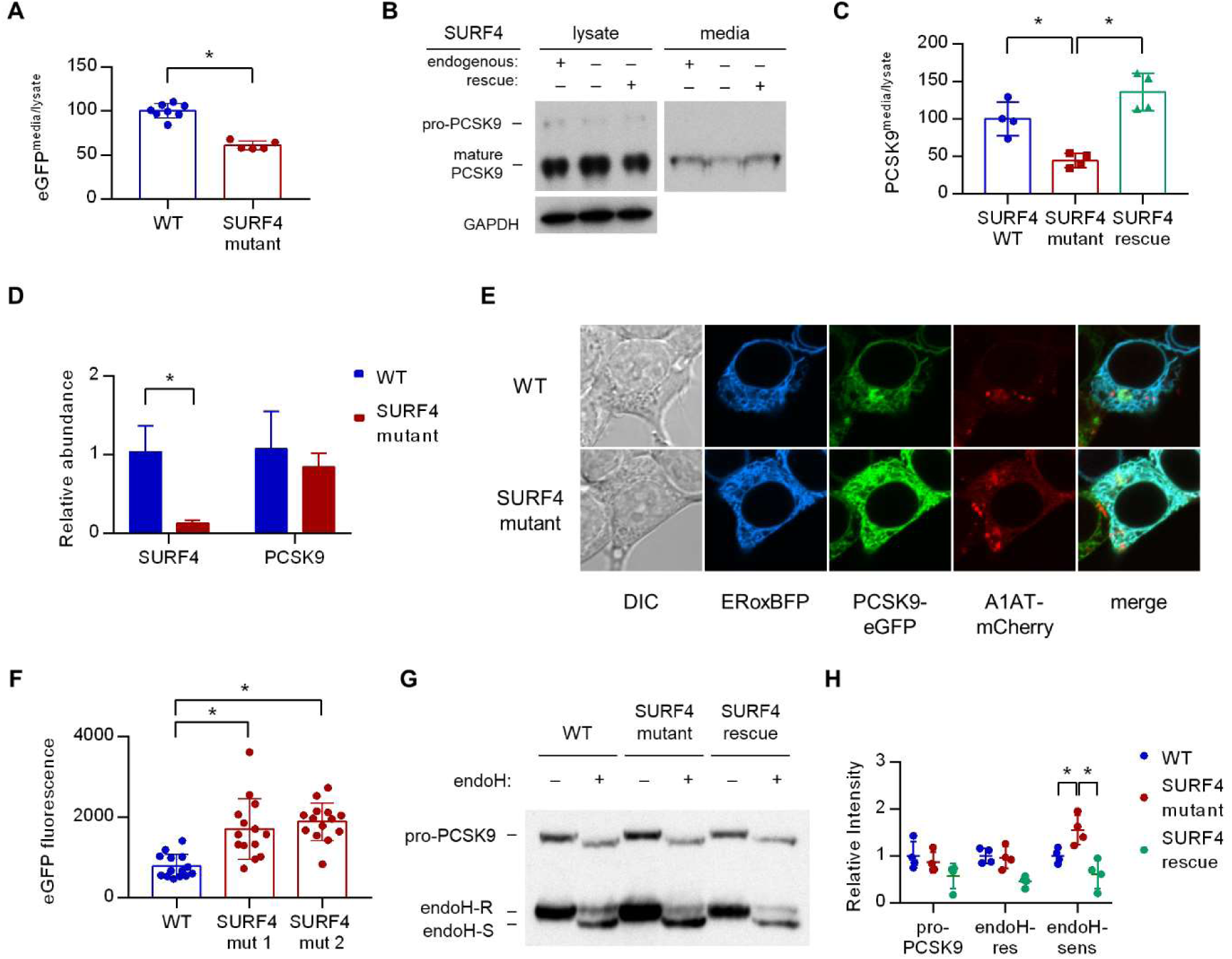
*SURF4* mutagenesis causes a decrease in PCSK9 extracellular secretion and an accumulation of PCSK9 in the ER. (A) Fluorescence detection of PCSK9-eGFP in extracellular conditioned media relative to cellular lysate in WT and clonal *SURF4*-deficient fluorescent reporter cell lines. (B) Immunoblotting of tetracycline-inducible PCSK9 in extracellular conditioned media and cellular lysates from WT, *SURF4*-deficient, or *SURF4* rescued cells. (C) Densitometry of immunoblots from (B). (D) Quantitative PCR of *SURF4* and PCSK9 transcript levels from RNA isolated from a *SURF4* WT or mutant fluorescent reporter cell line. (E) Live cell fluorescence microscopy of fluorescent reporter cells, either WT or *SURF4*-deficient, transfected with the ER marker ERoxBFP. (F) Quantification of PCSK9-eGFP signal intensity in pixels positive for ERoxBFP fluorescence. (G) EndoH-sensitivity of PCSK9 expressed in WT, *SURF4*-deficient, or *SURF4* rescued cells. (H) Quantification of endoH-sensitivity.

To identify other potential cellular components also required for efficient PCSK9 secretion, we next examined the 21 genes in addition to *SURF4* which exhibited potentially significant enrichment scores (FDR≤10%) in the whole genome CRISPR screen. PCSK9-eGFP-2A-A1AT-mCherry reporter cells were transduced with individual lentiviral CRISPR constructs targeting each of these genes and analyzed by FACS. None of the sgRNA targeting these additional genes were found to result in a significant effect specifically on PCSK9-eGFP fluorescence (Supplemental Figure 5). Thus, within the statistical power of our screen, *SURF4* emerges as the single gene out of the ∼19,000 human genes targeted by the GeCKOv2 library^18^whose inactivation results in specific intracellular retention of PCSK9 in HEK293T cells.

### SURF4 localizes to the early secretory pathway where it physically interacts with PCSK9

HEK293T cells were engineered to stably express *SURF4* with an N-terminal FLAG epitope tag. FLAG-tagged SURF4 demonstrated similar rescue of PCSK9-eGFP fluorescence in *SURF4*-deficient cells compared to native, untagged *SURF4* (Figure 3E), demonstrating that this tag does not interfere with SURF4 function. Consistent with previous reports^20,21^and compatible with a role for *S*URF4 as an ER cargo receptor, immunofluorescence of FLAG-*SURF4* demonstrated colocalization with a marker of the ER and, to a lesser extent, the ERGIC compartment (Figure 4A-B). Co-immunoprecipitation of cell lysates prepared in the presence of the chemical crosslinker dithiobis(succinimidyl propionate) demonstrated a physical interaction between FLAG-tagged SURF4 and GFP-tagged PCSK9 (Figure 4D).

### Loss of SURF4 results in decreased PCSK9 secretion and ER accumulation of PCSK9

Fluorescence assays on both extracellular conditioned media and intracellular lysates prepared from both wild-type and *SURF4*-deficient cells demonstrated a significantly decreased ratio of extracellular to intracellular PCSK9-eGFP fluorescence (Figure 5A), consistent with a defect in extracellular secretion. We also tested the dependence of untagged PCSK9 secretion on SURF4. Immunoblotting of cellular lysates and conditioned media demonstrated a significant decrease in the ratio of extracellular to intracellular levels in *SURF4*-deficient cells expressing untagged PCSK9 (similar to the decrease observed for the PCSK9-eGFP fusion), with rescue by stable expression of wild-type *SURF4* cDNA (Figure 5B-C). Quantitative PCR demonstrated equivalent levels of PCSK9 mRNA levels in wild-type and *SURF4*-deficient cells (Figure 5D), excluding an indirect effect on PCSK9 transcription.

To characterize the compartmental localization of intracellular PCSK9, we performed live cell fluorescence microscopy on wild-type and *SURF4*-deficient cells. PCSK9-eGFP fluorescence demonstrated increased colocalization with an ER marker in *SURF4*-deficient cells, consistent with ER retention (Figure 5E-F). Similarly, the predominant form of native PCSK9 in *SURF4*-deficient cells was sensitive to endoglycosidase H (Figure 5G-H), consistent with its localization in the ER^22^. Collectively these results indicate that *SURF4* promotes the efficient ER exit and secretion of PCSK9.

## DISCUSSION

It is estimated that ∼3000 human proteins are extracellularly secreted through the COPII pathway^23^, though few ER cargo receptors for these proteins have been identified and the proportion of secreted proteins that depend upon cargo receptor interactions is unknown^8^. Our findings suggest that SURF4 actively recruits PCSK9 into COPII vesicles, providing additional support for the cargo capture model of protein secretion. *SURF4*-deleted cells exhibit only a partial defect in PCSK9 secretion and whether the residual *SURF4*-independent secretion is due to bulk flow, or interactions with alternative ER cargo receptors, remains unknown.

To potentially circumvent the expected transient and/or low affinity nature of interactions between cargoes and their receptors, we applied purification by BirA*-mediated proximity-dependent biotinylation, which converts transient protein-protein interactions into permanent modifications^14^. Although this approach led to the detection of *SURF4* interactions with PCSK9, SAR1A, and SAR1B, *SURF4* was similarly marked by A1AT. Likewise, the A1AT cargo receptor LMAN1 was marked by both A1AT and PCSK9. These data suggest that BirA*-mediated proximity-dependent biotinylation may efficiently label ER cargo receptors, but lack the spatiotemporal resolution to distinguish incorporation within the same COPII vesicle from specific cargo – cargo receptor interactions.

CRISPR/Cas9-mediated gene editing has emerged as an efficient, programmable, and high-throughput tool for forward genetic screening in mammalian cells^24^. Our discovery of the PCSK9-SURF4 interaction demonstrates the feasibility of identifying ER cargo receptors by functional genomic screening and suggests that pooled CRISPR screening may prove to be a generalizable strategy applicable to the thousands of other secreted proteins for which a cargo receptor has not yet been identified. A limitation of our approach is its reliance on an immortalized cell line with the potential for accumulated somatic genetic and epigenetic changes, as well as restriction to a single cell-type gene expression program. Though LMAN1 cargoes appear to retain their cargo receptor-dependence for secretion in a variety of distantly related cell types^16,25,26^, the same many not apply to *SURF4*. Of note, SEC24A and/or SEC24B were previously shown to be required for efficient PCSK9 secretion in mice^9^. Though these genes were not identified in the current screen, this observation could be explained by the 5-10 fold greater expression of SEC24A than SEC24B in mouse liver, in contrast to the similar expression of both paralogs in HEK293T cells^27^.

In addition to a cargo receptor facilitating ER exit, our screening strategy should also survey other segments of the secretory pathway. Though sortilin has previously been implicated in post-Golgi transport of PCSK9^28^, sortilin dependence was not confirmed in a subsequent study^29^and also was not detected in the current screen. However, our data do not exclude a broader role for sortilin that also affected A1AT secretion, consistent with its contribution to the secretion of other proteins, including gamma-interferon^30^and ApoB-100^31^.

The yeast homologue of SURF4, Erv29p, was originally identified by proteomic analysis of purified COPII vesicles^32^. Erv29p promotes the secretion and concentrative sorting of the yeast mating factor gpαf into COPII vesicles through interaction with a hydrophobic I-L-V signal in gpαf ^19,33,34^. In HeLa cells, SURF4 has been shown to cycle in the early secretory pathway and to interact with LMAN1 and members of the p24 family. RNAi-mediated knockdown of *SURF4* alone resulted in no overt phenotype, though when combined with knockdown of LMAN1 caused morphologic changes to the ERGIC and Golgi compartments^20^. SURF4 has also been shown to interact with STIM1 and modulate store-operated calcium entry, though the mechanism underlying this observation is unclear^35^. Our screen did not identify significant enrichment of sgRNAs targeting p24 proteins, LMAN1, or STIM1, suggesting that these interactions are not required for efficient PCSK9 secretion in HEK293T cells.

Taken together with the marked reductions in plasma cholesterol associated with genetic or therapy-induced reductions in plasma PCSK9^3-6^, our findings raise the possibility that *SURF4* could represent an additional novel therapeutic target for the treatment of hypercholesterolemia. However, *SURF4* has not been identified in genome-wide association studies for human lipid phenotypes^36,37^, suggesting that partial reduction of *SURF4* expression does not limit PCSK9 secretion, consistent with the normal PCSK9 secretion and cholesterol profiles of Sec24A^+/-^ mice^9^and the normal levels of LMAN1 cargoes (coagulation factors V, VIII, and A1AT) in LMAN1^+/-^mice^17^. Though a loss-of-function variant (p.Gln185Ter) is present in ∼1:500 individuals^38^, no human diseases have been associated with *SURF4* deficiency and no mouse models for *SURF4* deletion have been reported^39^. Erv29, the *SURF4* homolog in yeast, is required for gpαf secretion and the C. elegans homolog (SFT-4) facilitates the secretion of yolk lipoproteins^21^, with the latter report also demonstrating a role for *SURF4* in apolipoprotein B secretion in HepG2 cells. Thus, it seems highly likely that additional *SURF4* cargoes remain to be identified.

## MATERIALS AND METHODS

### Cells and reagents

HEK293T cells were purchased from ATCC (Manassas VA). T-REx-293 cells were purchased from Invitrogen. Cells were cultured in DMEM (Invitrogen, Carlsbad CA) containing 10% FBS (D10) in a humidified 37°C chamber with 5% CO_2_. The expression construct for PCSK9-eGFP-2A-A1AT-mCherry was generated by Gibson assembly^40^of vector sequence derived from pNLF-C1 (Promega, Madison WI) and PCSK9 and A1AT cDNA derived by RT-PCR from HepG2 mRNA. Expression constructs for PCSK9, A1AT, SAR1A, and SAR1B fused to BirA* were generated by cDNA ligation into the entry vector pENTR/D-TOPO (Invitrogen) and Gateway cloning into the destination vector pDEST-pcDNA5-BirA-FLAG C-term^41^using LR clonase II (Invitrogen). This vector was also used as a backbone for the Gibson assembly of tetracycline-inducible expression of native PCSK9 and PCSK9-eGFP. For CRISPR experiments, sgRNA sequences were ligated into pLentiCRISPRv2 (Addgene #52961, a gift from Feng Zhang^18^) or pX459 (Addgene #62988, a gift from Feng Zhang) using BsmBI or BbsI restriction enzyme sites, respectively. Transfections were performed with FugeneHD (Promega) or Lipofectamine 3000 (Invitrogen) per manufacturer‘s instructions. Where indicated, clonal cell lines were derived by diluting cell suspensions to a single cell per well and expanding individual wells. Genotyping of clonal cell lines was performed by Sanger sequencing of target site PCR amplicons of genomic DNA isolated by QuickExtract (Epicentre, Madison WI). The pLentiCRISPRv2 whole genome CRISPR library (Addgene #1000000048, a gift from Feng Zhang^18^) was expanded by 8 separate electroporations for each half library into Endura electrocompetent cells (Lucigen, Middleton WI), plated on 24.5 cm bioassay plates, and pooled plasmids isolated by HiSpeed Maxi Prep (Qiagen, Hilden Germany). The pooled lentiviral library was prepared by co-transfecting 120 μg of each half library together with 120 μg of pCMV-VSV-G (Addgene #8454, a gift from Bob Weinberg^42^) and 180 μg psPAX2 (Addgene #12260, a gift from Didier Trono) into a total of 12 T225 tissue culture flasks of ∼70% confluent HEK293T cells using FugeneHD per manufacturer‘s instructions. Media was replaced at 12 hr post-transfection with D10 supplemented with 1% BSA, which was collected and changed at 24, 36, and 48 hr. Harvested media was centrifuged at 1000g for 10 min, pooled and filtered through a 0.45 μm filter, aliquoted, snap-frozen with liquid nitrogen and stored at -80°C until the time of use.

### Whole genome CRISPR screen

For each of 4 independent biological replicates, a total of ∼90 million cells stably expressing PCSK9-eGFP-2A-A1AT-mCherry were transduced at a multiplicity of infection of ∼0.3 with the whole genome CRISPR library. Puromycin selection (1 μg/mL) was applied from day 1 to day 4 post-transduction. Cells were passaged every 2-3 days and maintained in logarithmic phase of growth. After 14 days, a total of ∼240 million cells were detached with TrypLE Express (Invitrogen), harvested in D10 at 4°C, collected by centrifugation at 500g for 5 min, the pellet resuspended in 4°C phosphate-buffered saline (PBS) and filtered through a 35 μm nylon mesh into flow cytometry tubes, which were kept on ice until the time of sorting. A BD FacsAriaII was used to isolate cell populations containing ∼7 million cells per subpopulation into tubes containing D10. Genomic DNA was isolated using a DNEasy purification kit (Qiagen). Integrated lentiviral sgRNA sequences were then amplified using Herculase II polymerase (Agilent, Santa Clara CA) in a 2 step PCR reaction as previously described^18,43^with 20 cycles for round 1 PCR and 14 cycles for round 2. Amplicons were then sequenced on a HiSeq Rapid Run (Illumina, San Diego CA), with 95.2% of clusters passing quality filtering to generate a total of ∼210 million reads with a mean quality score of 34.99. Individual sgRNA sequences were mapped with a custom Perl script that seeded sequences onto a constant 24 nucleotide region upstream of the variable 20mer sgRNA, allowing up to 1 nucleotide mismatch, and reading the upstream 6 nucleotide barcode and downstream 20 nucleotide sgRNA sequence, which was then mapped to the library reference database with no mismatches tolerated. Enrichment was assessed using DESeq2 for individual sgRNA sequences^44^and MAGeCK for gene-level computations^45^.

### PCSK9 secretion assays

Clonal cell lines either wild-type for *SURF4* or containing frameshift-causing *SURF4* indels were isolated on a background of the HEK293T fluorescent reporter cell line or in T-REx-293 cells with a Flp/FRT-integrated PCSK9 cDNA. Cells were seeded at equal density in 10 cm plates and cultured in D10 for non-fluorescence-based assays or Fluorobrite media (ThermoFisher, Waltham MA) supplemented with 10% FBS for fluorescence-based assays. At the time of analysis, conditioned media was removed, clarified by centrifugation for 10 min at 1000g, and supernatant analyzed immediately or stored at -20°C. Cell monolayers were detached with trypLE express, collected in fresh media, pelleted, washed with PBS, and resuspended in 750 μL RIPA buffer (ThermoFisher) supplemented with protease inhibitors (Roche, Basel Switzerland). RIPA lysates were briefly sonicated, rotated end-over-end for 45 min at 4°C, and centrifuged at ∼20,000g for 30 min. Supernatants were transferred to a new tube and analyzed immediately or stored at -20°C. For fluorescence assays, samples were measured in triplicate in 96 well plates using an EnSpire fluorescence plate reader (PerkinElmer, Waltham MA), with fluorescent intensity zeroed on the autofluorescence of parental cells not expressing fluorescent fusion proteins for lysates or unconditioned media for conditioned media. For immunoblotting, samples were probed with antibodies against PCSK9 (Cayman 10007185, 1:1000), GAPDH (Abcam, Cambridge UK, ab181602, 1:10,000), GFP (Abcam, ab290, 1:5000), or (Abcam, ab167453, 1:1000). Densitometry was performed with ImageJ software^46^.

### Endoglycosidase H assays

To test for the N-glycosylation state of PCSK9, approximately 100 μg of RIPA lysate was incubated with denaturation buffer (NEB, Ipswich MA) for 10 minutes at 95°C, then split in half and incubated with or without 0.5 μL of PNGase (NEB) or EndoH (NEB) for 1 hr at 37°C.Laemmli sample buffer^47^was added, samples boiled for 5 min, resolved on a 10% Tris-HCl polyacrylamide gel, and analyzed by immunoblotting as above.

### Microscopy

Cells were grown on 35 mm poly-D lysine-coated glass bottom dishes (MatTek, Ashland MA). For live cell microscopy, cells were transiently transfected with an expression plasmid for ERoxBFP^48^(Addgene #68126, a gift from Erik Snapp^48^) and visualized at 24-48 hr post-transfection. For immunostaining, cells were washed with PBS, fixed with 2% paraformaldehyde, permeabilized with 0.1% Triton X-100 in PBS, blocked with 1% BSA and 0.1% Tween-20 in PBS, stained with FITC-conjugated anti-FLAG antibody (Sigma, St. Louis MO, F4049) and unconjugated rabbit antibodies against either calnexin (Cell Signaling Technology, Danvers MA, #2679), LMAN1 (Abcam, ab125006), or GM130 (Abcam, ab52649), then stained with Alexa647-conjugated anti-rabbit secondary antibody (Abcam, ab150075). All fluorescent imaging was performed on a Nikon A2 confocal microscope. Colocalization quantification was performed with Nikon Elements software. For all microscopy experiments, the observer was blinded to cell genotype and only unblinded after completion of quantitative analysis.

### BioID and mass spectrometry

BioID and mass spectrometry analysis was performed essentially as described^49^. Briefly, stable HEK293 Flp-In T-REx cells were grown on 15 cm plates to approximately 75% confluence. Bait expression and proximity labeling were induced by the addition of tetracycline (1 μg/mL) and biotin (50 μM) and proceeded for 24 hours. Cells were collected in PBS and biotinylated proteins purified by streptavidin-agarose affinity purification. Proteins were digested on-bead with sequencing-grade trypsin in 50 mM ammonium bicarbonate pH 8.5. Peptides were acidified by the addition of formic acid (2% (v/v) final) and dried by vacuum centrifugation. Dried peptides were suspended in 5% (v/v) formic acid and analysed on a TripleTOF 6600 mass spectrometer (SCIEX) in-line with a nanoflow electrospray ion source and nano-HPLC system. Raw data were searched and analyzed within ProHits LIMS^50^and peptides matched to genes to determine prey spectral counts^51^. High confidence proximity interactions (BFDR≤1%) were determined through SAINT analysis^52^implemented within ProHits. Bait samples (biological duplicates) were compared against 14 independent negative control samples (2 BirA*-FLAG-GFP only, 6 BirA*-FLAG only, and 6 3xFLAG only expressing cell lines) which were “compressed” to 6 virtual controls to increase the stringency in scoring^53^. Data has been deposited as a complete submission to the MassIVE repository (https://massive.ucsd.edu/ProteoSAFe/static/massive.jsp) and assigned the accession number MSV000082222. The ProteomeXchange accession is PXD009368.

### Mass spectrometry data analysis

All raw (WIFF and WIFF.SCAN) files were saved in our local interaction proteomics LIMS, ProHits^50^. mzXML files were generated from raw files using the ProteoWizard (v3.0.4468) and SCIEX converter (v1.3 beta) converters, implemented within ProHits. The searched database contained the human complement of the RefSeq protein database (version 57) complemented with SV40 large T-antigen sequence, protein tags, and common contaminants (72,226 sequences searched including decoy sequences). mzXML files were searched by Mascot (v2.3.02) and Comet (v2016.01 rev. 2) with up to two missed trypsin cleavage sites allowed and methionine oxidation and asparagine/glutamine deamidation set as variable modifications. The fragment mass tolerance was 0.15 Da and the mass window for the precursor was ±30 ppm with charges of 2+ to 4+ (both monoisotopic mass). Search engine results were analyzed using the Trans-Proteomic Pipeline (TPP v4.6 OCCUPY rev 3 check)^54^via iProphet^55^. Peptides with PeptideProphet scores ≥ 0.85 were mapped back to genes (gene IDs were from RefSeq). If peptides were shared between multiple genes, spectral counts were assigned exclusively to those genes with unique peptide assignments proportionally to the evidence for that assignment. If peptides matched only to genes without unique peptide assignments, spectral counts were divided equally between those genes^51^. SAINTexpress (v3.6.1)^52^was used to calculate the probability that identified proteins were significantly enriched above background contaminants using spectral counting (semi-supervised clustering) through comparing bait runs to negative control runs.

### Immunoprecipitation

Cells were harvested and resuspended at a density of ∼5×10^6^cells/mL in PBS supplemented with 2 mM CaCl2 and 2 mM dithiobis(succinimidyl propionate) (Pierce, Waltham MA), rotated end-over-end at 4°C, quenched with the addition of Tris-HCl (pH 7.5) to a final concentration of 25 mM, pelleted, and resuspended in lysis buffer (50 mM Tris-HCl, 150 mM NaCl, 2 mM CaCl2, and 1.0% Triton X-100, supplemented with protease inhibitors (Roche), pH 7.5). Lysates were prepared as described above. Immunoprecipitation from lysates was performed with M2-FLAG affinity gel (Sigma) or GFP-trap magnetic beads (Chromo-Tek, Hauppage NY). To reduce nonspecific binding, M2-FLAG affinity gel was pre-blocked with 1 hr incubation in High Salt Wash Buffer (50 mM Tris-HCl, 500 mM NaCl, 2 mM CaCl2, pH 7.5) supplemented with 5% BSA. Pulldowns were performed with 500 µL of lysate with end-over-end rotation at 4°C overnight. A total of 5 washes were performed with Wash Buffer (50 mM Tris-HCl, 500 mM NaCl, 2 mM CaCl2, pH 7.5) for GFP-trap beads or High Salt Wash Buffer for M2-FLAG affinity gel. Protein elution was performed by incubating beads at room temperature for 15 min with 2X Laemmli sample buffer supplemented with β–mercaptoethanol.

### Screen validation

All genes identified by MaGECK with a FDR<10% were selected for follow up validation. The top-ranking sgRNA for each gene was individually cloned into BsmBI sites of pLentiCRISPRv2. Individual lentiviral stocks were prepared and used to transduce fluorescent reporter cells at an MOI <0.5, followed by puromycin selection, and passaging for 2 weeks prior to FACS analysis. The mean fluorescence intensity of a total of 20,000 gated events was recorded for each construct and compared to the mean intensity of 3 nontargeting sgRNA constructs. A total of 3 biologic replicates was performed.

### Statistical Analysis

The statistical significance of differences in quantitative data between control and experimental groups was calculated using the student‘s *t*-test. CRISPR screen and mass spectrometry data analysis was performed as described above.

## Acknowledgments

This research was supported by NIH grants R35-HL135793T (D.G.) and T32-HL007853 (B.T.E.). D.G. is a Howard Hughes Medical Institute investigator.

## Author contributions

B.T.E. and D.G. conceived the study, designed experiments, and wrote the manuscript. B.T.E., G.G.E., E.K., V.T., P.J.L., and J.X. performed experiments. All authors contributed to data analysis and manuscript review.

---

**Supplemental Figure 1.**
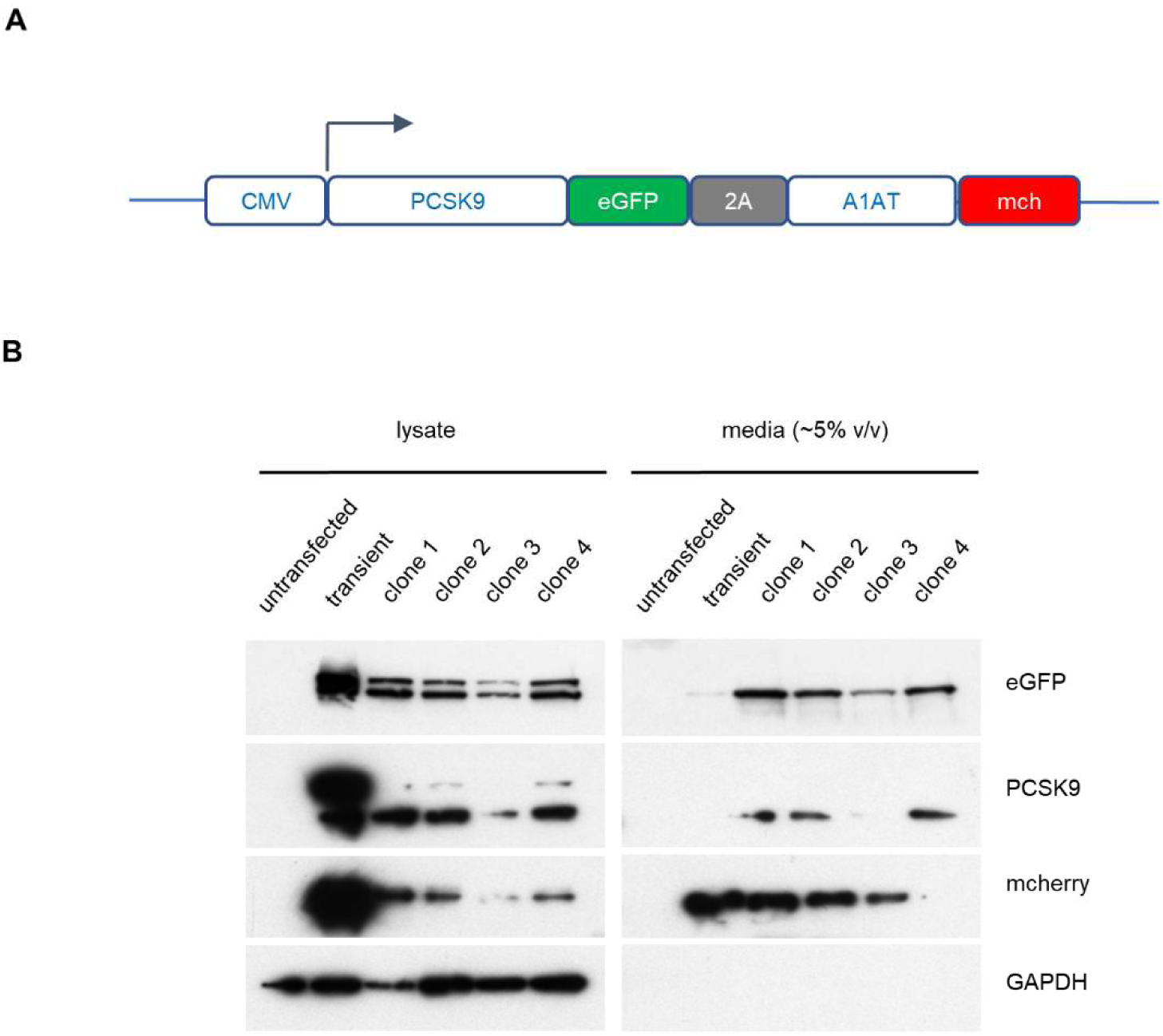
Analysis of PCSK9-eGFP-2A-A1AT-mCherry reporter cell clones. (A) Construct for CMV-promoter-driven expression of PCSK9-eGFP-2A-A1AT-mCherry expression. (B) Immunoblotting of HEK293T cells transfected with the PCSK9-eGFP-2A-A1AT-mCherry vector, and 4 individual drug-resistant clonal cell lines, relative to parental untransfected cells.

**Supplemental Figure 2.**
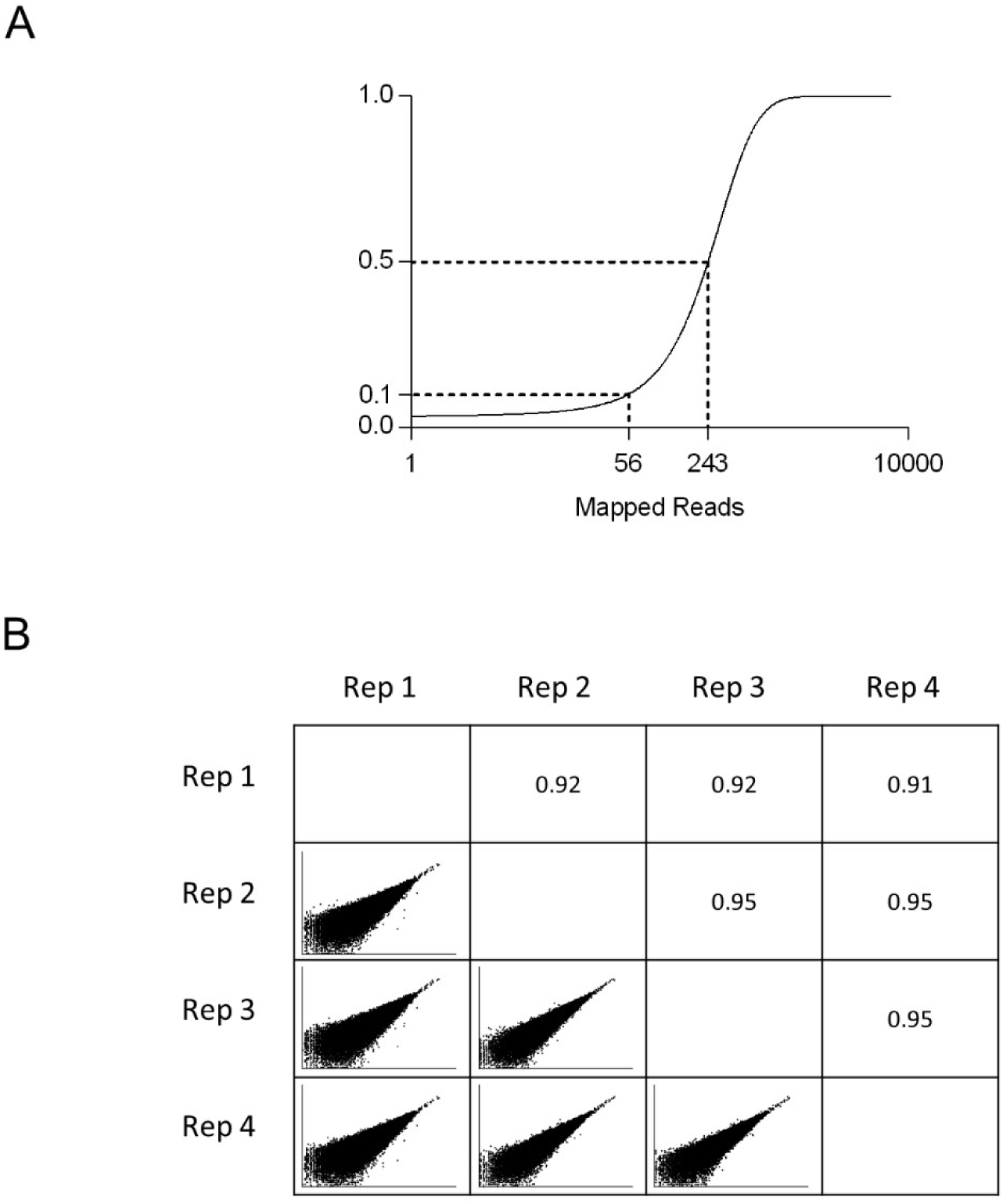
Whole genome screen analysis. (A) Cumulative distribution function of mapped reads for each sgRNA in the whole genome library. (B) Comparison of read counts for every pairwise combination of 4 biologic replicates with Pearson correlation coefficient.

**Supplemental Figure 3.**
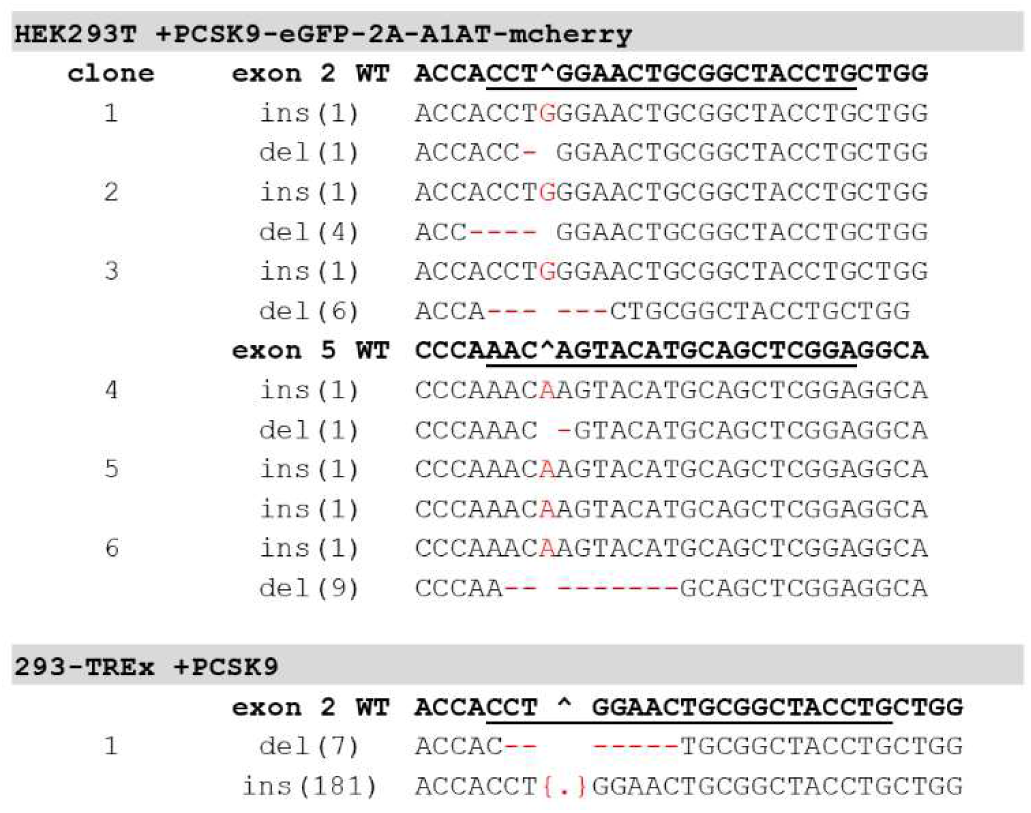
*SURF4*-deficient genotypes. Clonal cell lines were genotyped by PCR amplification of sgRNA target site and Sanger sequencing of the amplicon with either TIDE decomposition of individual alleles from the amplicon, or ligation of the amplicon into cloning plasmids and Sanger sequencing of multiple individual clones.

**Supplemental Figure 4.**
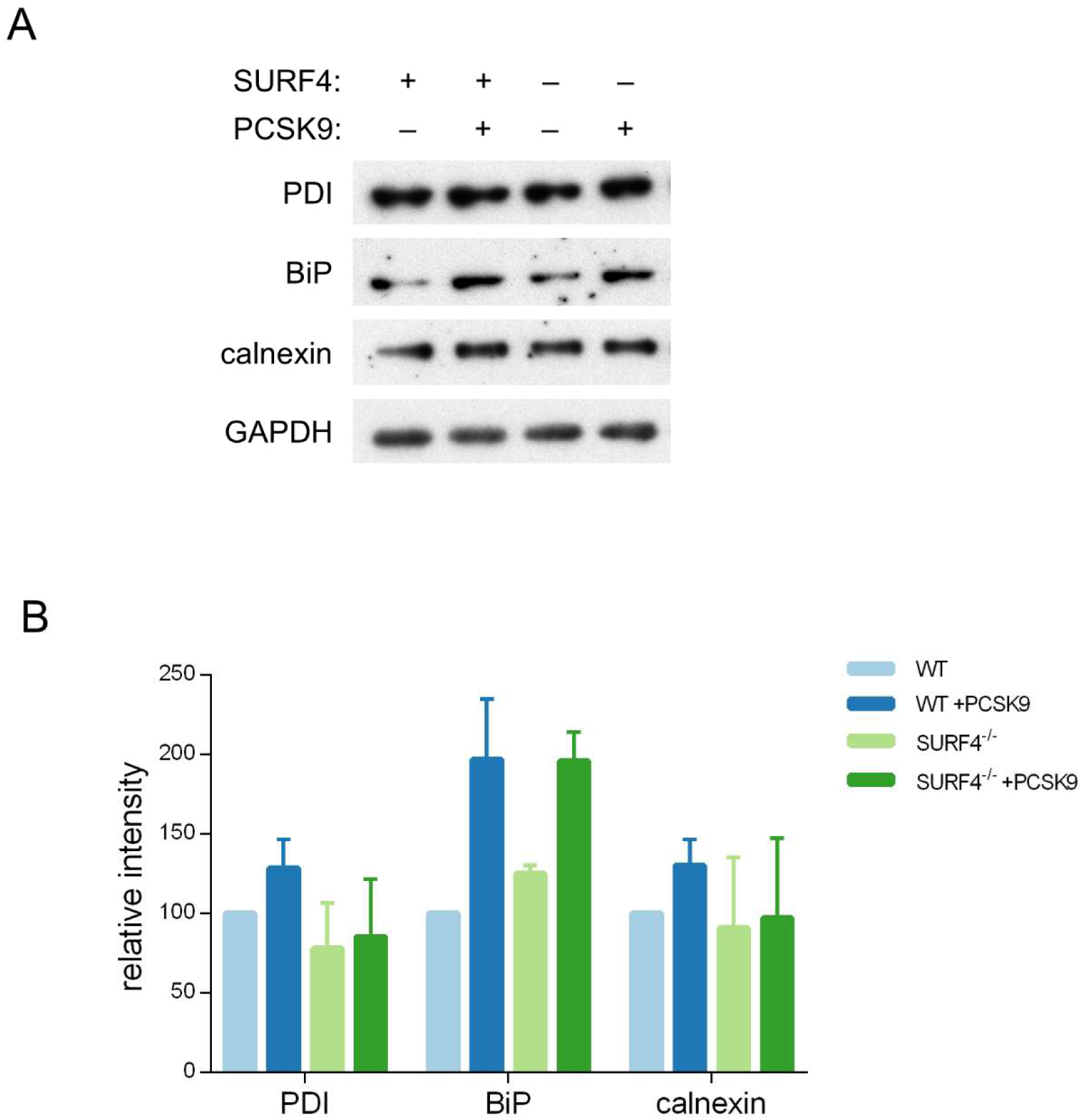
ER stress markers. Clonal cell lines with a stably integrated tetracycline-inducible PCSK9 cDNA on either WT or *SURF4* mutant background were treated with tetracycline or vehicle control and analyzed by immunoblotting for various markers of ER stress (A), quantified by densitometry (B).

**Supplemental Figure 5.**
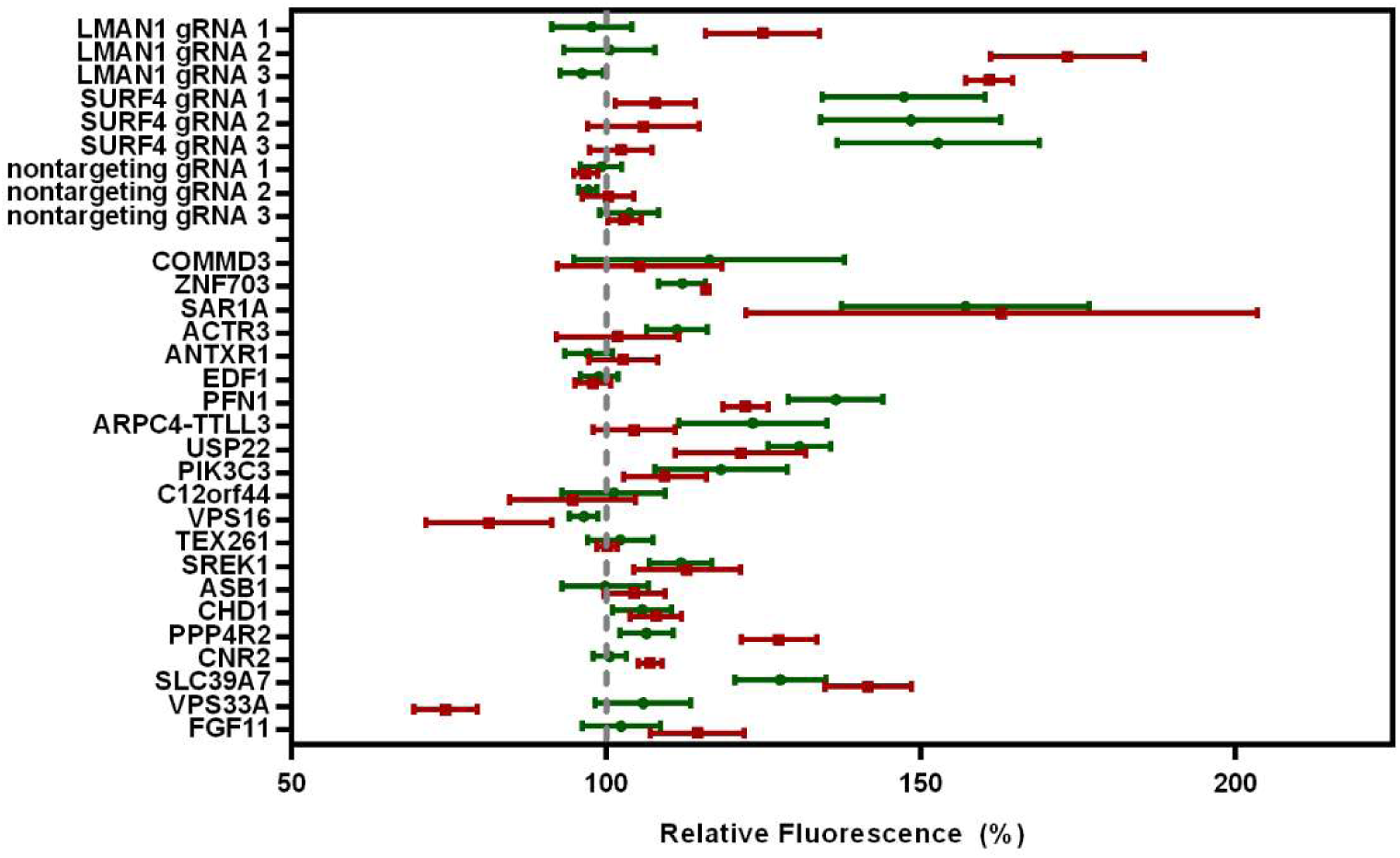
Validation experiments for additional candidate genes. The top-scoring sgRNA for each of the next 21 most enriched genes after *SURF4* was individually cloned into lentiCRISPRv2 and tested in PCSK9-eGFP-2A-A1AT-mCherry reporter cells. Flow cytometry was performed for each of 3 biologic replicates. The mean fluorescent intensity of PCSK9-eGFP and A1AT-mCherry for sgRNA against each candidate gene relative to nontargeting sgRNA controls is shown.

**Supplemental Table 1. BioID of PCSK9-interacting proteins.** Spectral counts are listed for prey proteins identified from streptavidin-purified lysates of cells expressing fusions of BirA* with GFP, PCSK9, A1AT, SAR1A, and SAR1B. Gene Ontology enrichment analysis^56,57^is displayed for candidate PCSK9 cargo receptors.

**Supplemental Table 2. CRISPR screen sgRNA-level analysis**. DESeq2 output for each sgRNA showing normalized mapped reads in control (GFP-low) populations and log2 fold-change in experiment (GFP-high) populations with significance testing for enrichment or depletion. A copy of the reference library showing the corresponding gene target and target sequence and gene for each sgRNA is included.

**Supplemental Table 3. CRISPR screen gene-level analysis**. Enrichment for each sgRNA targeting the same gene was combined to generate a gene-level enrichment score using MAGeCK^45^, which is displayed in the column corresponding to the number of unique sgRNA targeting that same that demonstrated enrichment (p<0.05) by analysis with DESeq2 (Supplemental Table 2).

**Supplemental Tables 1-3:** Uploaded as separate files given size limitations.

**Supplemental Table 4.**
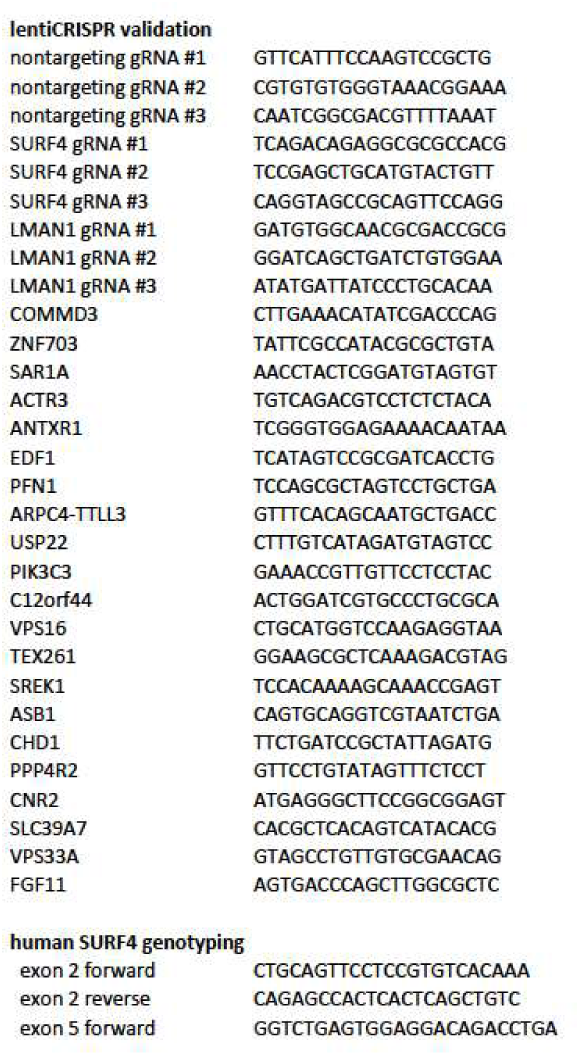
Oligonucleotide sequences. Primers used for CRISPR/Cas9 targeting and genotyping are displayed.

